# Neural bases of phonological and semantic processing in early childhood

**DOI:** 10.1101/858613

**Authors:** Avantika Mathur, Douglas Schultz, Yingying Wang

## Abstract

During the early period of reading development, children gain phonological (letter-to-sound mapping) and semantic knowledge (storage and retrieval of word meaning). Their reading ability changes rapidly, accompanied by their learning-induced brain plasticity as they learn to read. This study aims to identify the specialization of phonological and semantic processing in early childhood using a combination of univariate and multivariate pattern analysis. Nineteen typically developing children between the age of five to seven performed visual word-level phonological (rhyming) and semantic (related meaning) judgment tasks during functional magnetic resonance imaging (fMRI) scans. Our multivariate analysis showed that young children with good reading ability have already recruited the left hemispheric regions in the brain for phonological processing, including the inferior frontal gyrus (IFG), superior and middle temporal gyrus, and fusiform gyrus. Additionally, our multivariate results suggested that the sub-regions of the left IFG were specialized for different tasks. Our results suggest the left lateralization of fronto-temporal regions for phonological processing and bilateral activations of parietal regions for semantic processing during early childhood. Our findings indicate that the neural bases of reading have already begun to be shaped in early childhood for typically developing children, which can be used as a control baseline for comparison of children at-risk for reading difficulties.

## Introduction

In today’s society, learning to read as a child is the foremost step for developing high literacy skills. Teaching a child to read begins at birth with the reinforcement of pre-literacy skills, and most children officially learn to read between the ages of 5 and 7 years old. Two common approaches to teach reading are sounding-out and sight-word reading methods. The sounding-out approach asks children to read aloud and to pronounce each letter or group of letters to recognize words by their sounds, which helps children to build letter-to-sound knowledge. Meanwhile, the sigh-word approach requires children to memorize sight words or common vocabulary, which allows children to build their internal lexical dictionary. Thus, word reading can be achieved through grapho-phonological processing and lexico-semantic processing. The neural bases of these two processes have been studied mainly in older children and adults (Coltheart et al., 2001; Jobard et al., 2003; Vigneau et al., 2011), but not in young children (5 – 7 years old). To understand the neural bases of early reading is critical for not only providing evidence on theoretical models of reading development, but also building control baseline to be used for examining how children with reading difficulties differ.

The two dominate theoretical models of reading are the dual-route cascaded (DRC) model (Coltheart et al., 2001) and the parallel-distributed-processing (PDP) connectionist model (Harm and Seidenberg, 2004). The DRC model suggests two distinct routes for word reading including the grapho-phonological route transforming visual words into their sound representations (indirect route) and the lexico-semantic route transforming visual words into their meanings (direct route). According to the DRC model, skilled adult readers identify family words and words with irregular pronunciations like ‘pint’ (via the direct route) and pronounce newly encountered words and nonwords by rule-based grapheme-to-phoneme mapping (via the indirect route). In contrast, the PDP model postulates a single mechanism that generates pronunciations for all words. The PDP model suggested that word pronunciations are learned through repeated training with a corpus of written and spoken inputs. Different neural pathways have been identified to support the DRC model of reading (Friederici and Gierhan, 2013; Hickok and Poeppel, 2007; Price, 2012). A systematic meta-analysis of the DRC model of reading in adults’ brain has proposed a dorsal route for grapho-phonological processing and a ventral route for lexico-semantic processing (Jobard et al., 2003). The dorsal route consists of the left superior temporal gyrus (STG), the left dorsal inferior parietal lobe (IPL, covering supramarginal gyrus and angular gyrus), and the left opercular part of the inferior frontal gyrus (opIFG). The ventral route is composed of the left fusiform gyrus (FG), the left basal inferior temporal area, the left posterior part of the middle temporal gyrus (MTG), and the left orbital part of IFG (orIFG). Additionally, the triangular part of the left IFG (trIFG) has been suggested to be recruited in both routes. For PDP model, words are pronounced using the same neural networks after a set of optimal connection weights has been learned (Binder et al., 2005; Harm and Seidenberg, 2004). The PDP model produces outputs stimulating children’s learning behavior. The DRC model suggests that the young readers relie more on the indirect grapho-phonological route to translate letters into corresponding sounds and less on the direct lexico-semantic route to derive a meaningful representation of a given word. With practice over time, the young readers develop a larger internal lexicon dictionary storing words that are can be recognized by sight via the lexical processing without the semantic processing.

The current literature on the neural representations of phonological and semantic processing in typically developing children is limited to late childhood (8 – 15 years old) (Bitan et al., 2007a; Bitan et al., 2007b; Blumenfeld et al., 2006; Cao et al., 2009; Hoeft et al., 2007; Li et al., 2018). Atypical brain structures and functions have been identified in children at-risk for developmental dyslexia in early childhood (5 – 7 years old) (Im et al., 2016; Raschle et al., 2012; Raschle et al., 2014; Wang et al., 2016; Yu et al., 2018) and even as early as in infancy (Langer et al., 2015; Sideridis et al., 2019). Thus, there is limited knowledge of the neural bases of reading in typically developing children during early childhood when they began to learn to read. To study the neural bases of reading will provide more evidence to improve our understanding of the DRC model and might help us to identify children at-risk for reading difficulties during early childhood. Early identification can lead to early intervention that has been shown to be more effective at the time when brain plasticity is high.

FMRI studies have showed brain activation in the bilateral FG, the left STG/MTG, the left SPL, the bilateral precentral gyrus, and the left IFG/SFG/MFG using English-word rhyming judgment tasks (Bitan et al., 2007a; Cao et al., 2006; Cao et al., 2009; Hoeft et al., 2006; Hoeft et al., 2007). FMRI studies have identified the neural bases of semantic processing in the bilateral FG, ITG/MTG, IPL, and IFG using English-word semantic judgement tasks (Blumenfeld et al., 2006; Booth et al., 2001; Booth et al., 2003; Booth et al., 2004). Few studies have directly compared brain activation for phonological and semantic processing in young children using visual word pairs. One fMRI study directly compared brain activation between phonological and semantic processing and found greater brain activation in the bilateral FG and MFG/IFG comparing semantic processing versus phonological processing in 26 adolescents with a wide age range (9 – 19 years old) (Landi et al., 2010). However, other studies reported that there was no significant difference between phonological processing and semantic processing in Chinese children (9 – 13 years old) (Cao et al., 2009; Liu et al., 2018). Their findings suggested that the neural representations of reading in children have not yet been specialized to the DRC model of reading. However, reading Chinese words is quite different from reading English words, which might contribute to the disparate findings. A recent longitudinal fMRI study provided neural evidence for development of the DRC model of reading in older children (8 – 14 years old) (Younger et al., 2017). They reported that phonological decoding is initially used with gradually decreasing reliance as reading becomes more automated for a child. Their fMRI results indicated that increases in reading ability were associated with decreases in brain connectivity of the dorsal route (indirect grapho-phonological route) and increases in brain connectivity of the ventral route (direct lexico-semantic route). In summary, past research suggested that the neural bases of reading supporting the DRC model have not yet specialized until adulthood and the specialization of neural pathways is directly associated with individual reading ability.

The advancement of fMRI data analysis led to the application of powerful pattern-classification algorithms to examine multi-voxel patterns of brain activation. The multi-voxel pattern analysis (MVPA) has been suggested to more sensitive and flexible for examining cognitive states (Norman et al., 2006). The conventional fMRI analysis uses subtraction-based approach to identify brain activation related to certain experimental versus control conditions at a voxel-by-voxel level, which might overlook the brain activations if differences between conditions are across multiple voxels instead of a single voxel (Norman et al., 2006). To achieve greater sensitivity for discriminating conditions of interest with greater power and flexibility than the conventional univariate analysis (Kriegeskorte et al., 2006), we used searchlight analysis to identify differences of neural patterns across different conditions.

The present study aimed to examine the neural bases of phonological and semantic processing in nineteen young children (5 – 7 years old) using fMRI during child-friendly visual rhyming and semantic judgement tasks during early childhood. We hypothesized that bilateral frontal and temporo-parietal brain regions would be involved in both phonological and semantic processing. The use of both univariate and MVPA analyses will provide important complementary information (Poldrack and Farah, 2015). Thus, we conducted both univariate analysis and MVPA to understand the neural bases of phonological and semantic processing during early childhood.

## Materials and Methods

### Participants

60 healthy native English-speaking children were recruited for the study. The study involved two visits. During the first visit, a series of standardized behavioral tests were administered. The participant’s parent was asked to fill out a questionnaire about the child’s developmental history. Based on the questionnaire, participants with 1) a diagnosis of Attention Deficit Hyperactivity Disorder (ADHD), 2) hearing or vision impairment, 3) neurological or psychiatric disorders, 4) a diagnosis of language disorder or reading disability, or 5) with any contraindications to be scanned in an MRI, were excluded from the study and not invited for the second visit. The inclusion criteria for the second visit (imaging session) were: (1) native monolingual English-speakers, (2) righthanded as measured by handedness questionnaire (Oldfield, 1971), (3) above 80 standard score of the non-verbal intelligence measured by the Kaufman Brief Intelligence Test, Second Edition (Kaufman, 2004), (4) accuracy greater than 70% in the familiarity task. The computer-based familiarity task was designed to ask the participant to read thirty 3-5 letter monosyllabic words and choose the corresponding picture that represents the word. The purpose of this customized picture naming task is to make sure that the participant would be able to complete both rhyming and semantic judgement tasks during fMRI scans. 20 participants were invited back for the fMRI session based on the inclusion criteria mentioned above. FMRI data from one participant were excluded due to excessive motion artifacts. 19 participants were included in the final analysis for the present study. They were all right-handed (8M, 11F; mean age of 6.55 years old with age range from 5 year 4 months – 7 year 9 months). The Institutional Review Board approved all experimental procedures at the University of Nebraska at Lincoln. Written consent forms were obtained from the parent or guardian and written assent forms were obtained from children who were older than seven years old.

### Behavioral Measures

Behavioral measures were acquired during the first visit. Phonological awareness (PA) was measured using the PA subtest of WRMT-III (WRMT-III-PA) (Woodcock, 2011), which consists of five sections including first-sound matching, last-sound matching, rhyme production, blending, and deletion tasks. Word reading ability was measured using the word identification subtest of WRMT-III (WRMT-III-WID) (Woodcock, 2011). Word association skill was measured by the word classes sub-test of the core language score in the CELF-5 (Wiig et al., 2013). The CELF score was only administered for 12 participants.

### Imaging Data Acquisition

#### Paradigms

Three-to five-letter monosyllabic words suitable for young children were selected from the MRC (Medical Research Council) psycholinguistic database (Wilson, 1988) to generate word-pair stimuli matched for concreteness, printed familiarity, word type (noun), and the number of syllables. Both rhyming and semantic judgment tasks presented word stimuli visually with a child-friendly image above the corresponding words (Figure 1). Each word was presented for a duration of 800 ms with a 200-ms gap between word-pair presentations. Then, a 2,200-ms response screen followed after the presentation of each word-pair stimulus, during which the word-pair stimulus remained on the screen along with a question mark in the middle of the two words (Figure 1). Participants were instructed to respond as soon as they saw the question mark. For the rhyming judgment task, the participant presses the right button (color red) if the two words rhyme and presses the left button (color blue) if the two words do not rhyme. For the semantic association judgement task, the participant presses the right button (color red) if the two words associate semantically and presses the left button (color blue) if they do not associate semantically. The control conditions are the same for both tasks and consisted of three to five non-alphabetic glyph characters of symbol strings. The two strings were also presented with child-friendly images to matched the task condition for controlling visual inputs. During the control condition, the participants were asked to determine whether the symbol strings were matched or not. In addition, there are 10-second baseline conditions consisted of a black cross on white screen. The participants were instructed to press either the left or right button as soon as they saw the black cross change to a red cross (Figure 1).

We administered two runs for each paradigm to minimize the effects of fatigue. A block design was used to achieve higher detectability of brain activation in pediatric fMRI data (Wilke et al., 2003). Each run consisted of eight blocks (four task blocks and four control blocks) starting with a fixation block (baseline) followed by a task block, a fixation block, and a control block (Figure 1D). The order of runs was counterbalanced across subjects, such that nine participants completed rhyming paradigm first, whereas the other 10 completed semantic paradigm first. Each rhyming block consisted of eight conditions (four rhyming/four non-rhyming). Each semantic block comprised eight conditions (four semantic association/four non-semantic association). Each control block consisted of four conditions (two matching strings/two non-matching strings). Within a block (either task or control block), stimuli were randomized. At the beginning of each run, 5-sec fixation was presented as a baseline block and were not included in the fMRI data analysis to eliminate non-equilibrium effects of magnetization. Each run lasted for 4 min 37 sec (277 Seconds). Each run consisted of 32 task conditions 16 control conditions, and 8 fixation conditions.

**Figure 1.**
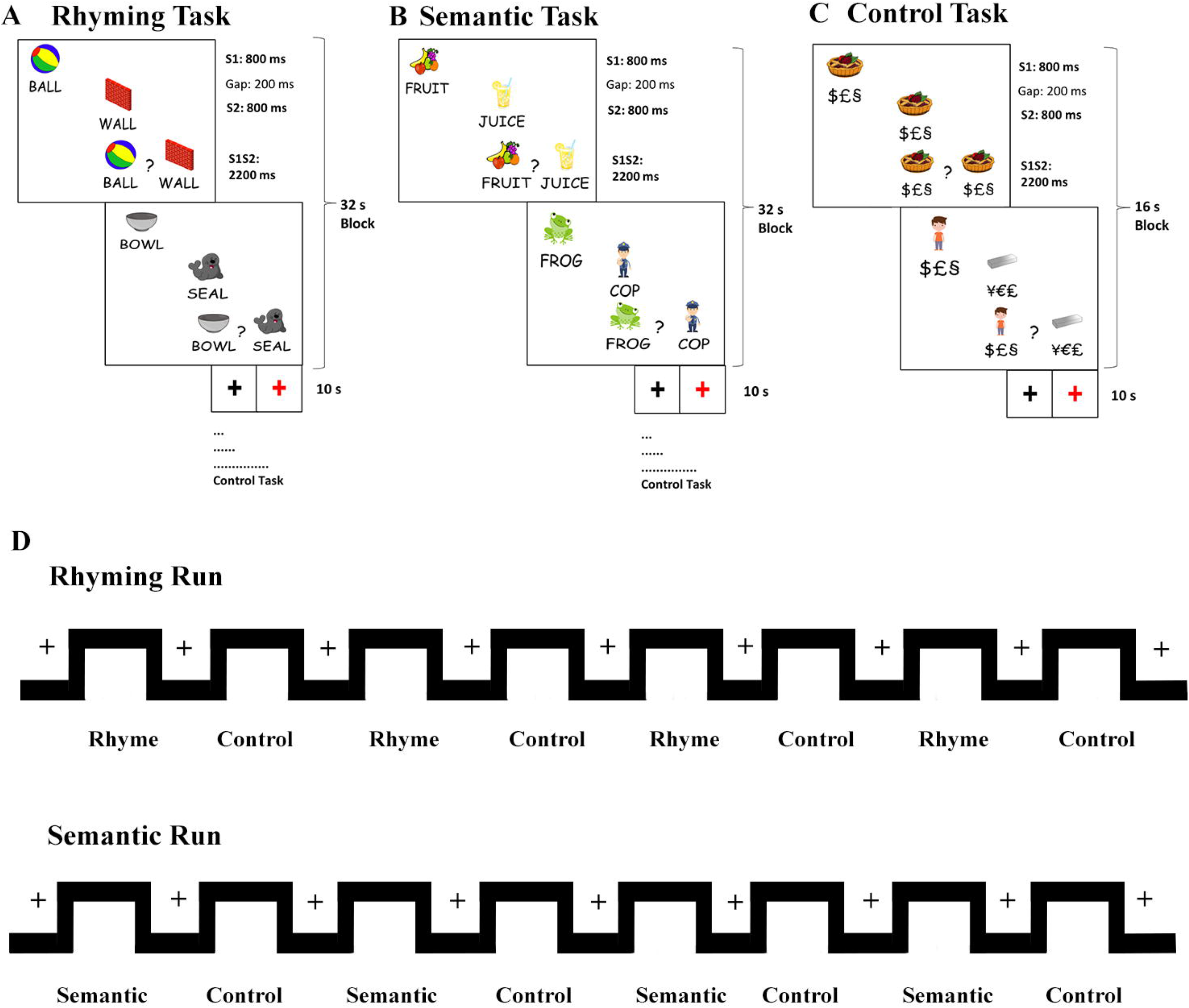
Schematic representation of fMRI task design.

### Imaging Acquisition Protocol

Brain imaging data were acquired using a 3.0-Tesla Skyra Siemens scanner with a 64-channel head coil. The blood-oxygen-level-dependent (BOLD) signal was measured using a multiband EPI (University of Minnesota sequence cmrr_mbep2d_bold) sequence with the following parameters: TR = 1,000 ms, TE = 29.80 ms, flip angle = 60°, matrix size = 210×210 mm^2^, field of view = 210 mm, slice thickness = 2.5 mm, and number of slices = 51. Voxel size = 2.5×2.5×2.5 mm^3^. Before functional image acquisition, a high resolution T1-weighted 3D structural image was acquired for using a multi-echo magnetization prepared rapid gradient echo (MPRAGE) sequence (TR = 2,530 ms, TE1 = 1.69 ms, TE2 = 3.55 ms, TE3 = 5.41 ms, TE4 = 7.27 ms, matrix size = 256×256 mm^2^, field of view = 256 mm, slice thickness = 1 mm, number of slices = 176, and TA = 6.03 minutes).

Prior to the imaging session, each participant underwent 30-min practice training in a child-friendly “MRI-like” room (a mock scanner room equipped with a nonmagnetic MRI simulator, Psychology Software Tools, Model#100355). The literature suggests that pre-training sessions are crucial for young children and increases the success rates, and reduce the motion artifacts and anxiety (Leach and Holland, 2010). We achieved a 95% success rate by using a mock session. During the mock session, participants were exposed to different scanner noise and practiced the experimental tasks until they achieved accuracy above 60%. The practice version of the experimental tasks used words that were different to the actual stimuli. During the actual imaging session, each participant laid on the scanning table with his/her head secured with foam pads. A two-button box was placed on each side of the participant and responses were collected through E-prime 2 (Psychology Software Tools, Inc.). We attached elastic straps across the head-coil apparatus to reduce the head motions. The state-of-art OptoActive™ active noise canceling headphones (OptoAcoustics, Mazor, Israel) was used to minimize the effects of the ambient scanner noise.

### Imaging Data Analyses

#### Univariate Analysis

Standard preprocessing steps were performed in SPM12 (Wellcome Department of Cognitive Neurology, London, https://www.fil.ion.ucl.ac.uk/spm/). Re-alignment and ArtRepair (Mazaika et al., 2007) were used to correct for head motions. The parameters for Art-Repair outlier detections were: global mean signal change z = ± 9 and exceeding 4 mm of head motion. Outliers and volumes with excessive motion identified by ArtRepair were de-weighted in the generalized linear model (GLM) analysis. Structural MRI data were segmented and normalized to the segmented pediatric template NIHPD (4.5-8.5 years) based on brain imaging data from 82 healthy young children obtained from the Neuroimaging and Surgical Technologies Lab, MNI, Canada (http://nist.mni.mcgill.ca/) (Fonov et al., 2011). Then, functional images were coregistered to normalized structural images. Finally, fMRI data were smoothed using an 8×8×8 mm^3^ isotropic Gaussian kernel. Four conditions (Fixation, Rhyming, Semantic, and Control) were modeled using the first-level GLM framework and each run was modeled in a separate GLM. Head motion and outlier were regressed out in the GLM analysis. A high-pass filter with a 128-s cutoff and an artificial mask threshold of 0.2 were applied. Contrasts were then defined to reveal brain regions specifically involved in the phonological processing (Rhyming > Control) and semantic processing (Semantic > Control). The first-level statistical parametric maps (SPMs) were further entered into a second-level random-effects analysis. Significant clusters were identified at a cluster threshold k > 20 with FDR correction (q < 0.05). The in-scanner accuracy scores were used as covariate of no interest to control individual variance due to task difficulty. A direct comparison between Rhyming > Control and Semantic > Control was performed to identify differences of brain activity between the phonological processing and the semantic processing. Additionally, a conjunction analysis between Rhyming > Control and Semantic > Control was performed to identify the brain regions that are commonly involved to both processes.

#### Multivariate Pattern Analysis

We employed a whole-brain searchlight analysis that is a recently developed MVPA technique for identifying locally informative areas of the brain. The searchlight analysis outperforms mass-univariate analyses due to its higher sensitivity to distributed information coding (Kriegeskorte et al., 2006). We performed searchlight analysis using a linear discriminant analysis (LDA) classifier implemented in CoSMoMVPA toolbox (Oosterhof et al., 2016). Two separate searchlight analyses were conducted to examine the spatial pattern of voxels in the brain that the classifier could reliably distinguish between (1) rhyme task from control task, and (2) semantic task from control task. The first-level SPMs from each run were stacked together into the searchlight analysis with 100 voxel searchlight spheres across the whole brain. Classification accuracies were obtained using a leave-one-out cross-validation method with an eight-fold partitioning scheme for each subject. For each run, the dataset was split into eight chunks (each corresponding to one experimental block), and the classifier was trained on the data from seven chunks and tested on the remaining one. The procedure was repeated for eight iterations, using all possible train/test partitions. The average decoding accuracies across these iterations were calculated. At the group level, we performed a two-tailed one-sample t-test across individual maps where classification was significantly above chance (50%, since our classifiers were binary). The resulting SPMs were corrected for multiple comparisons using a cluster-based Monte Carlo simulation algorithm with 1000 iterations implemented in the COSMOMVPA toolbox (corrected cluster threshold α = .01, two-tailed; z > 1.96).

## Results

### Behavioral Measures

The mean PA standard score of the 19 participants was 113.16 with standard deviation (SD) of 13.85. The mean standard score of WRMT-III-WID was 112.76 with SD of 14.41. The CELF word association subtest acquired for 12 participants had the mean scaled score of 12.83 with SD of 2.37. The in-scanner performance data of accuracy and reaction time (RT) (for corrected responses) were plotted in Figure 2 and summarized in Table 1. Children performed better in control conditions as compared to task conditions. A paired sample t-test revealed that the accuracy was significantly higher in the control task than the rhyming task (t (18) = −3.23, p <.05) and the semantic task (t (18) = −3.52, p <.05). RT was significantly lower in the control task than the rhyming task (t (18) = 3.24, p <.05) and the semantic task (t (18) = 2.91, p <.05). There are no significant differences in accuracy and RT between the rhyme and semantic judgment tasks.

**Figure 2.**
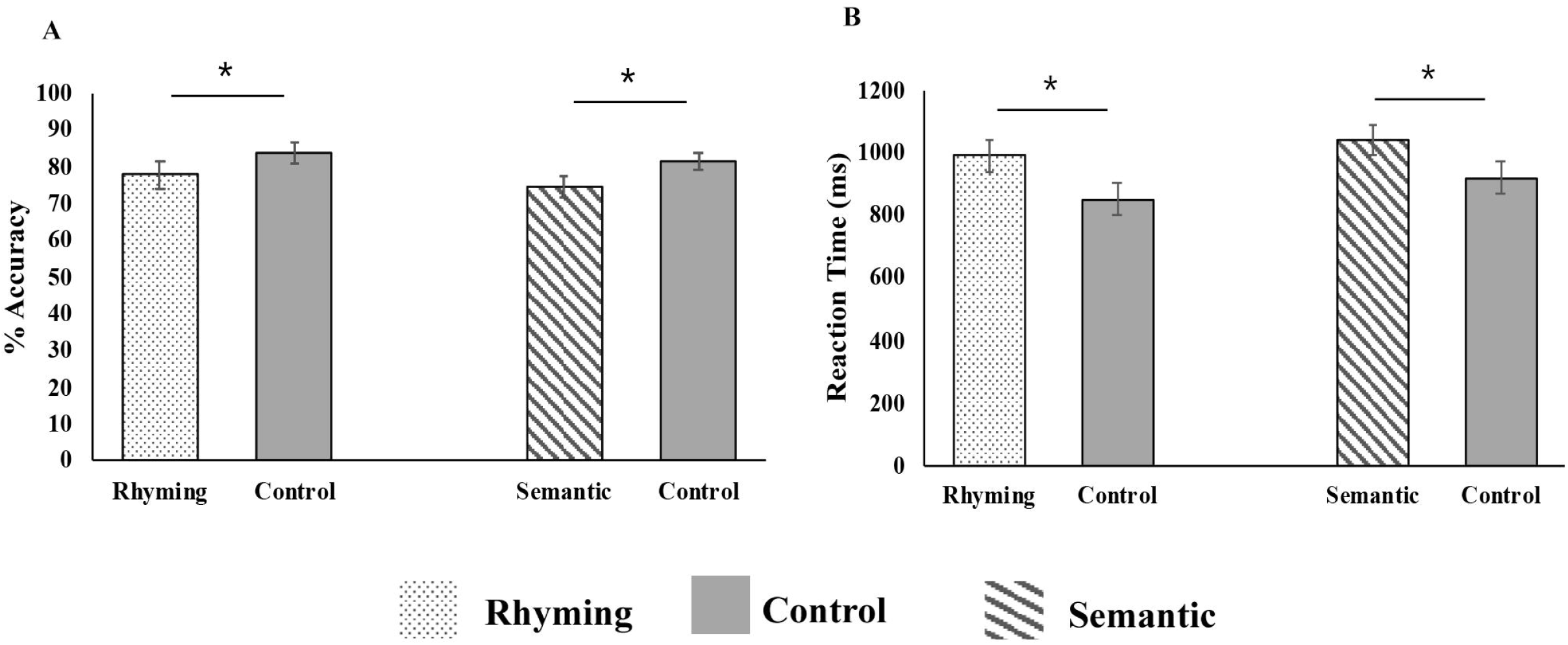
Average (N=19) in-scanner task accuracy and reaction time.

**Table 1.**
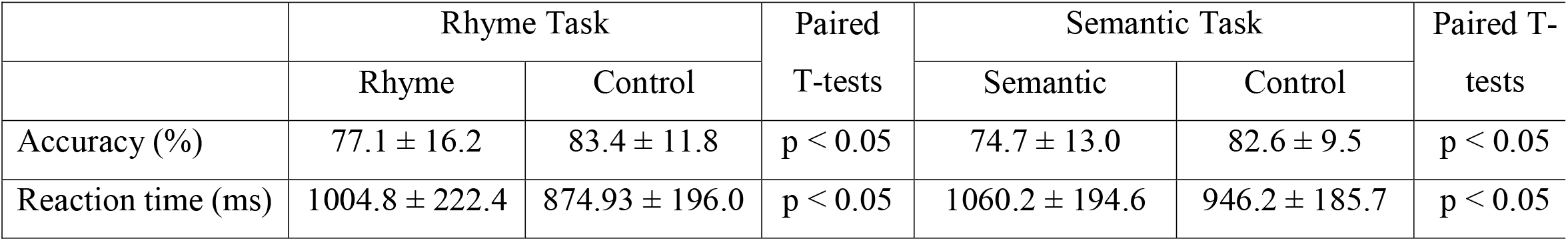
In-scanner Performance

### Univariate Analysis

The univariate results of rhyme and semantic judgement tasks were summarized in the Table 2 and 3 (Figure 3). The contrast of rhyme > control identified significant activations in the left frontal regions including the left IFG covering opIFG, trIFG, and orIFG, the left supplementary motor area (SMA), and the left precentral gyrus (Figure 3A and Table 2). The contrast of semantic > control showed significant activations in the left IFG (opIFG, trIFG, orIFG), which also extended to the right IFG (trIFG and orIFG), and the left precentral gyrus. There were significant activations in the bilateral temporo-parietal regions covering the right MTG/ITG and FG, as well as IPL including the supramarginal gyrus, angular gyrus and precuneus (Figure 3B and Table 3).

The direct comparison between the rhyming and semantic task did not show any significance of either direction (rhyming > semantic or rhyming < semantic) after FDR correction (q < 0.05) with a cluster threshold k > 20. The conjunction analysis of the two contrasts (rhyme versus control and semantic versus control) revealed overlap in activations in the left IFG and the left SMA after FDR correction (q < 0.05) with a cluster threshold k > 20 (supplementary Figure 2 and supplementary Table 1).

**Figure 3.**
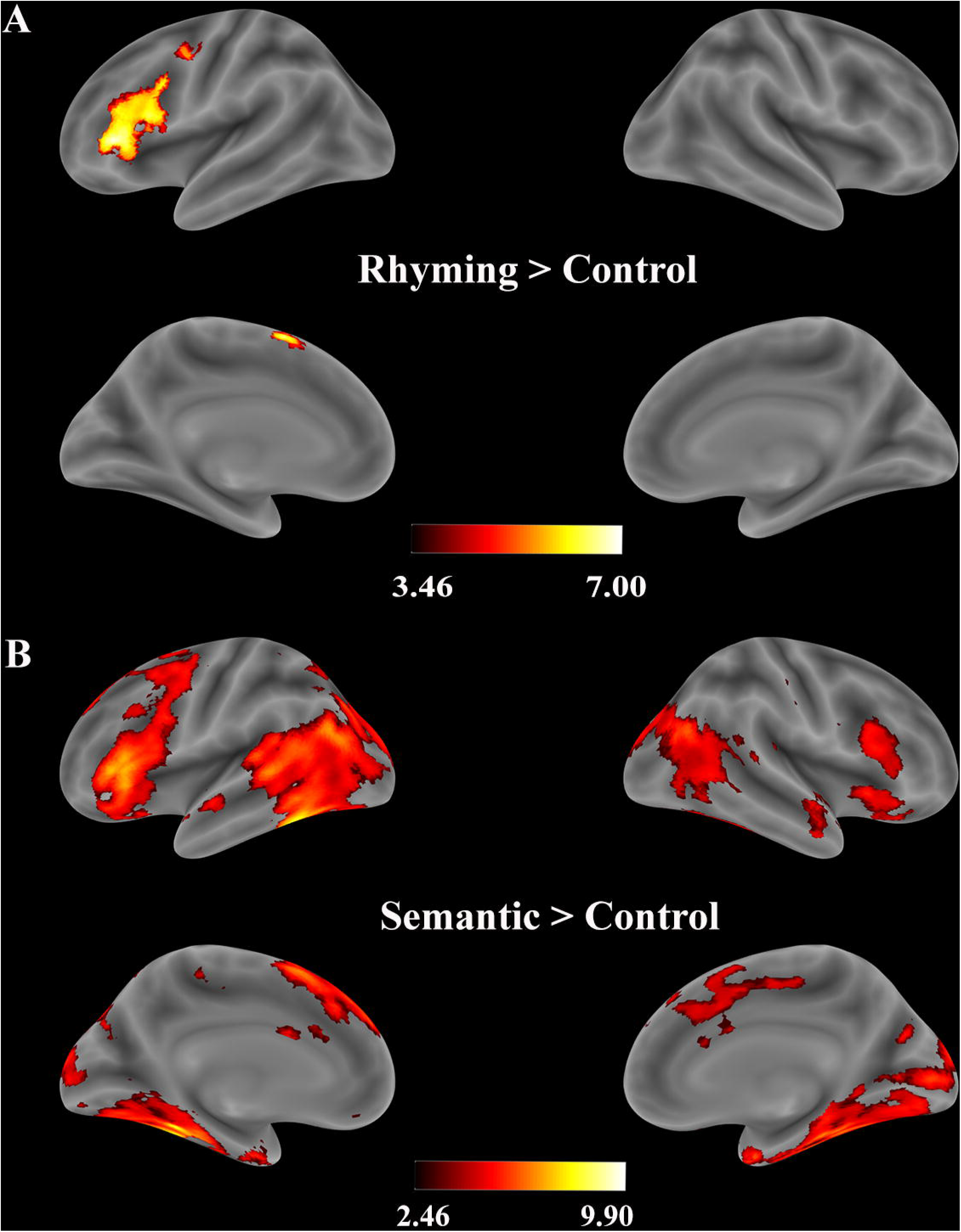
Voxel-wise significant activation, within the whole brain anatomical mask for the contrast (A) rhyming vs control and (B) semantic vs control. The color bar indicates test statistics (t-score) for the significant clusters identified using a voxel-wise threshold of q < 0.05, FDR correction at a cluster threshold k > 20.

**Table 2.**
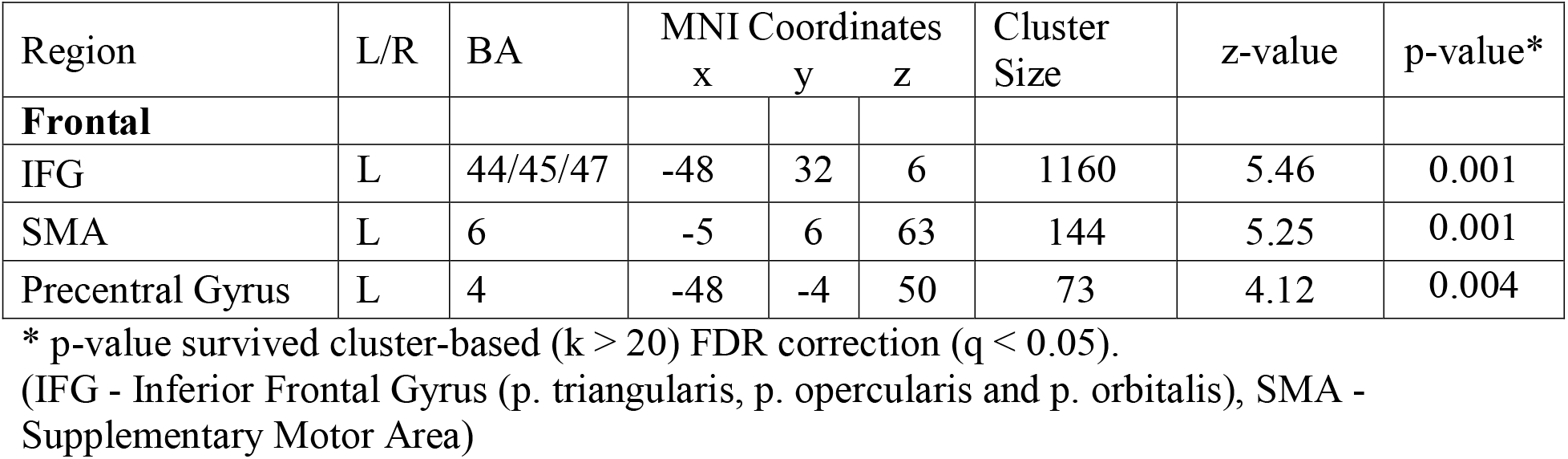
Univariate results for the contrast Rhyme vs Control.

| Region | L/R | BA | MNI Coordinates |  |  | Cluster Size | z-value | p-value* |
| --- | --- | --- | --- | --- | --- | --- | --- | --- |
|  |  |  | x | y | z |  |  |  |
| **Frontal** |  |  |  |  |  |  |  |  |
| IFG | L | 44/45/47 | -48 | 32 | 6 | 1160 | 5.46 | 0.001 |
| SMA | L | 6 | -5 | 6 | 63 | 144 | 5.25 | 0.001 |
| Precentral Gyrus | L | 4 | -48 | -4 | 50 | 73 | 4.12 | 0.004 |

\* p-value survived cluster-based ( $k > 20$ ) FDR correction ( $q < 0.05$ ).
(IFG - Inferior Frontal Gyrus (p. triangularis, p. opercularis and p. orbitalis), SMA - Supplementary Motor Area)

**Table 3.**
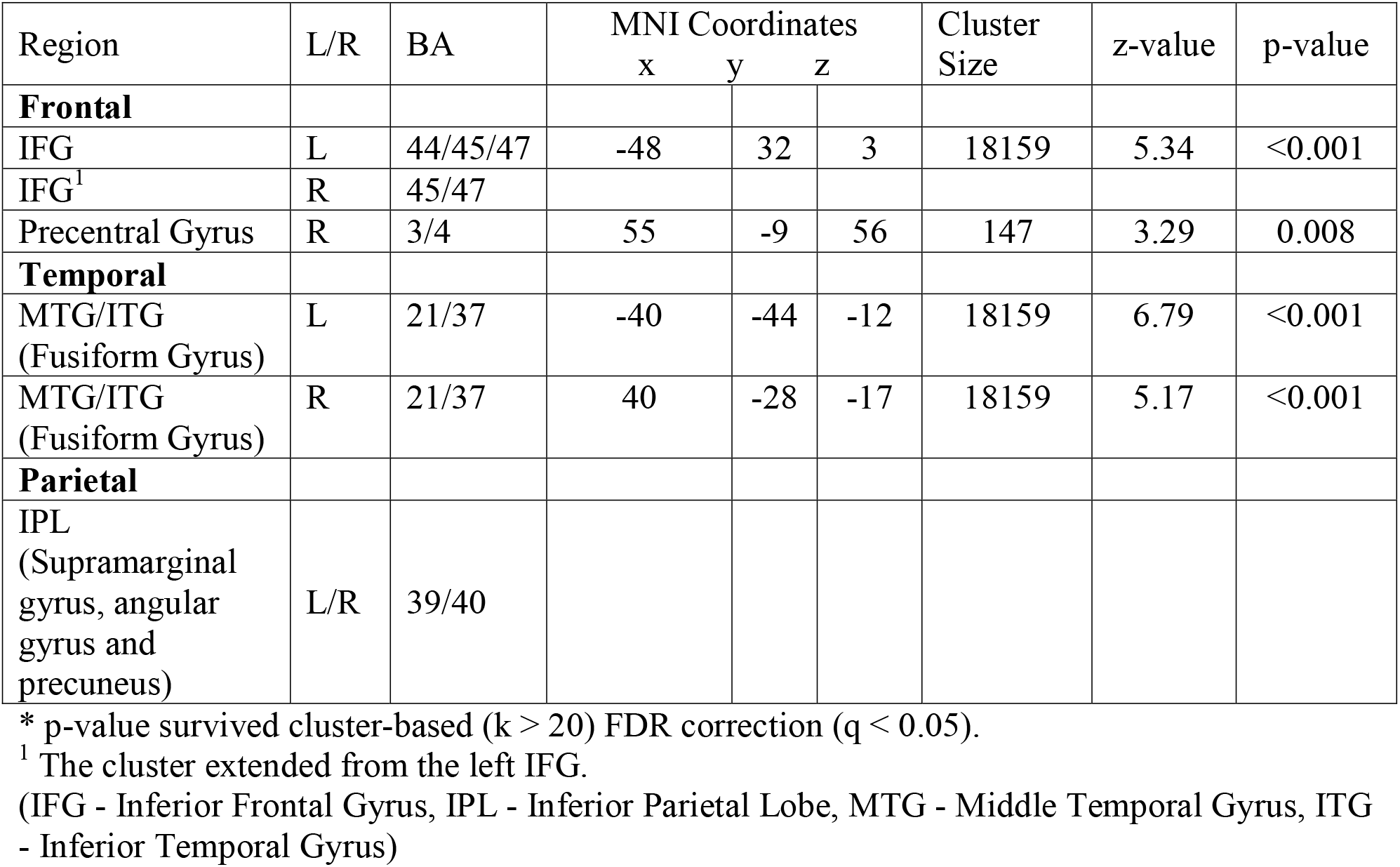
Univariate results for the contrast Semantic vs Control.

### Multivariate Analysis

For rhyme versus control, searchlight analysis identified significant decoding accuracy in the left fronto-temporal regions including the left opIFG and the left pre/postcentral gyrus, and the left MTG/STG/ITG and FG (Figure 4A and Table 4). For semantic versus control, searchlight analysis found significant decoding accuracy in the left fronto-temporal and bilateral parietal regions. Amongst the frontal regions, the left trIFG cluster extended to the left precentral gyrus, and the left MFG. Amongst the temporal regions, a cluster extending from the left STG/MTG to the left ITG (including FG) significantly decoded semantic conditions from control conditions. The temporal cluster also extended to the bilateral parietal regions (IPL: supramarginal gyrus, angular gyrus, and precuneus) (Figure 4B, Table 5).

**Figure 4.**
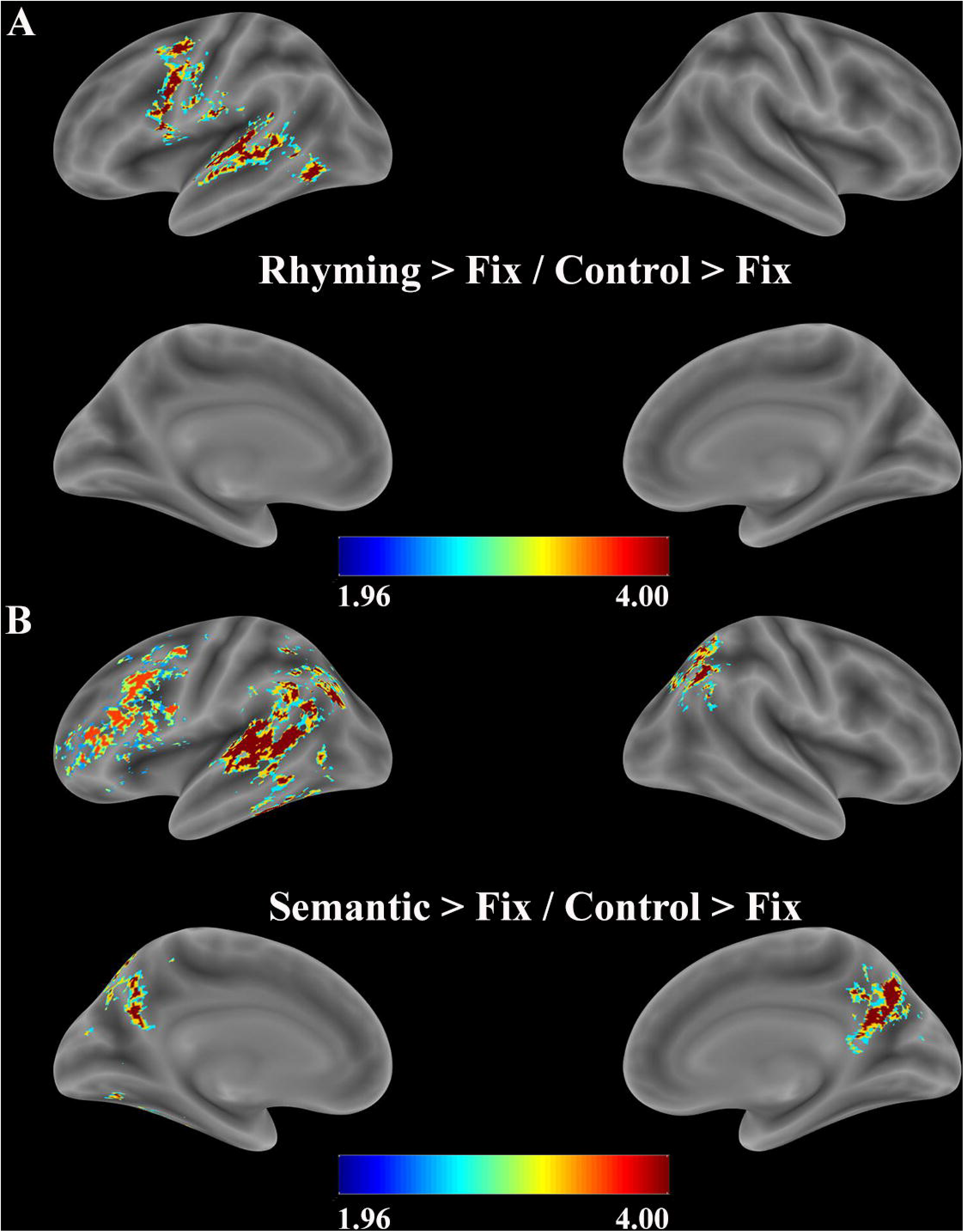
Multivariate Pattern analysis results. Statistical group maps for the two-searchlight analysis performed with a 100-voxel searchlight using LDA classifier to identify regions that significantly decode above chance (A) rhyme from control condition and (B) semantic from control condition. The resulting statistical maps were corrected for multiple comparisons using a cluster-based Monte Carlo simulation algorithm implemented in the COSMOMVPA toolbox [Oosterhof et al., 2016, clusterstat maxsum function] (corrected cluster threshold α = .01, twotailed; z > 1.96). The color bar represents the test statistics (z-score) for the clusters with significant decoding accuracy.

**Table 4.**
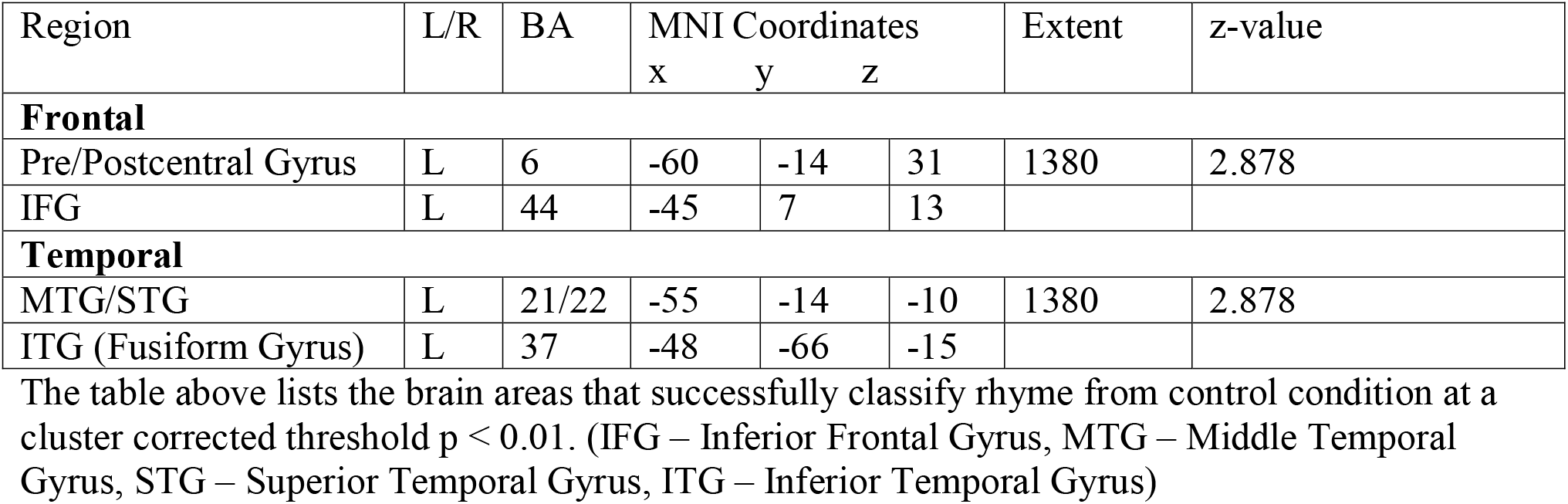
MVPA searchlight results for Rhyme vs Control.

**Table 5.** MVPA searchlight results for Semantic vs Control.

| Region | L/R | BA | MNI Coordinates |  |  | Extent | z-value |
| --- | --- | --- | --- | --- | --- | --- | --- |
|  |  |  | x | y | z |  |  |
| **Frontal** |  |  |  |  |  |  |  |
| IFG<sup>1</sup> | L | 45 | -35 | 29 | -15 | 1680 | 2.327 |
| **Temporal** |  |  |  |  |  |  |  |
| MTG/STG | L | 21/22 | -53 | -11 | 3 | 3762 | 2.878 |
| ITG (Fusiform gyrus) | L | 37 | -45 | -61 | -5 |  |  |
| **Parietal** |  |  |  |  |  |  |  |
| IPL (Supramarginal gyrus, angular gyrus and precuneus) | L/R | 39/40 |  |  |  |  |  |

The table above lists the brain areas that successfully classify semantic from control condition at a cluster corrected threshold $p < 0.01$ .
<sup>1</sup> This cluster also extends to the left precentral gyrus (BA 6) and the left MFG (BA 46). (IFG – Inferior Frontal Gyrus, MTG – Middle Temporal Gyrus, STG – Superior Temporal Gyrus, ITG – Inferior Temporal Gyrus, IPL – Inferior Parietal Lobe)

## Discussion

While performing the visual phonological judgment tasks, children have to access the sounds of the visually presented words and are therefore more likely to engage the grapho-phonological route of reading (Bitan et al., 2006; Bitan et al., 2007a; Bitan et al., 2007b; Booth et al., 2007; Cao et al., 2006; Cao et al., 2009; Hoeft et al., 2007). Semantic association tasks require children to decide if the two words presented are related and thus are more likely to engage the lexico-semantic route of reading (Blumenfeld et al., 2006; Booth et al., 2001; Booth et al., 2003; Booth et al., 2004). In this study, we examined the neural representations of early reading skills using visual rhyming and semantic judgement tasks during fMRI in young children (5 – 7 years old). Our univariate results indicated that the phonological processing in the brain had a left lateralized activation pattern, whereas the semantic processing in the brain involved a bilateral activation pattern. However, our MVAP results suggested both processes in the brain involved left-lateralized patterns. The present study provided evidence that young children with good reading ability have already established the grapho-phonological route for reading in the left hemisphere to support early stages of reading including the left IFG, the left STG/MTG and the left FG. In addition, our results suggested that the lexico-semantic processing also relies on the left IFG, the left STG/MTG and the left FG, as well as the bilateral parietal regions covering IPL. The MVPA results provided a detailed account of neural representations of IFG sub-regions and suggested that the left opIFG was specifically involved for phonological processing, whereas the left trIFG was specifically involved for semantic processing. Additionally, the bilateral parietal regions showed specialization for semantic processing. All these findings will be discussed in more detail below.

### Frontal Lobe

Our univariate results showed left lateralization in the IFG, SMA, and precentral gyrus for both tasks and no significant differences through direct comparison between tasks at FDR-corrected level. In line with a recent study on 35 typically developing children (5 – 6 years old) (Weiss et al., 2018), our direct comparison results supported that there is no specialization in the frontal regions yet during early childhood. However, our multivariate results indicated that phonological processing and semantic processing involved different sub-regions of the left IFG. The left opIFG was specialized for phonological processing, whereas the left trIFG was specialized for semantic processing (Table 4 and 5). In adults, it has been reported that the phonological processing relied on the left opIFG (Jobard et al., 2003). Moreover, real-time transcranial magnetic stimulations (TMS) to the left opIFG disrupted phonological processing (Gough et al., 2005), suggesting that the left opIFG as the posterior part of the left IFG was involved in phonological processing. Our MVPA results also supported that the left IFG is not a single functional region. The left trIFG as the ventral part of the left IFG was involved for tasks that focus on semantics (Price, 2012). Different from our univariate results, our MVPA findings indicated that young typically developing children have already show some specialization of the sub-regions of the left IFG for phonological and semantic processes by the age of 5 – 7 years. The differences of results between the univariate and multivariate analyses may be caused by the higher sensitivity and greater power of the MVPA approach (Kriegeskorte et al., 2006). The MVPA compares the representation patterns of activity across conditions, while the univariate analysis compares spatial-average activation across conditions.

Both univariate and multivariate results showed a left lateralization for the left IFG for phonological processing. Booth et al. used visual rhyming task in both adults and older children and only found activation in the right IFG in adults (Booth et al., 2004). Our findings on the left IFG are aligned with their results, suggesting that young children also recruited a left lateralized activation patterns in the IFG during visual rhyming task.

### Temporal Lobe

Previous research suggested that the left MTG was identified for semantic processing in both young (5 – 6 years old) and older children (9 – 12 years old) (Bitan et al., 2007b; Blumenfeld et al., 2006; Booth et al., 2004; Weiss et al., 2018). Studies that have directly compared phonological and semantic processing in adults’ brain have reported a double-dissociation between the tasks in the temporal regions (Jobard et al., 2003). In adults, the left STG was involved in the phonological processing, while the semantic processing recruited the left MTG (Binder, 2016; Devlin et al., 2003; McDermott et al., 2003; Poldrack et al., 1999; Price, 2012). Weiss et al. directly compared the early specialization of brain regions for phonological and semantic processing of spoken language during early childhood (5 – 6 years old) using auditory rhyme and semantic judgment tasks, respectively (Weiss et al., 2018). By comparing the differences of brain activation between the two tasks using subtraction-based univariate analysis, they found specialization of the left STG and the left supramarginal gyrus for phonological processing, and the left MTG for lexical processing. Their findings suggested that the temporal regions have been already specialized for spoken language by 5 years of age. However, our univariate results showed no FDR-corrected activation in temporal regions for phonological processing. The discrepancy might be due to different task stimuli. They used auditory presented words, whereas we used visually presented words.

The left lateralization for reading has been reported to be related to children’s reading ability in early childhood (Yamada et al., 2011). Yamada et al. used a one-back letter reading task on 5-year-old children who received reading instruction in kindergarten and found that typically developing children with on-track pre-literacy skills recruited the left-lateralized temporal regions in the brain. In contrast, children at-risk for reading difficulty showed more bilateral activation. Thus, the left hemispheric lateralization observed in our study could be a result of good pre-reading skills of the children recruited in our study.

### Temporo-occipital Lobe

The bilateral FG has been involved in processing visually presented words that requires orthographic representations in older children (9 – 12 years old) (Booth et al., 2004). A letter box also known as the visual word form area (VWFA) in the left FG has been suggested to decode letter strings to words (McCandliss et al., 2003). Turkeltaub et al. showed a decrease in activation in the right VWFA (anatomically homologous to the left VWFA) with an increase in age (6 – 22 years) (Turkeltaub et al., 2003). They concluded that learning to read led to decreased activations in the right temporo-occipital regions accompanied by increased activations in the left IFG/MTG. Previous research suggested that the left VWFA has already visible by the age of 7 years (Gaillard et al., 2003; Parviainen et al., 2006). Our results showed the involvement of the left VWFA in both visual rhyming task and semantic task and provided evidence that VWFA has already specialized to left hemisphere by early childhood (5 – 7 years of age). While some authors postulate that the VWFA is dedicated solely to the lexico-semantic route (Levy et al., 2009), others propose that VWFA is common for both grapho-phonological and lexico-semantic routes, and information is then be passed on to the most appropriate route for reading a word (Goswami, 2008; Jobard et al., 2003). A recent study showed the involvement of the left VWFA during a phonological awareness task in 5-6-year old children (Wang et al., 2018). Our findings supported that the left VWFA is recruited for both pre-reading routes during early childhood.

### Parietal Lobe

We identified bilateral parietal regions (supramarginal, angular gyrus, and precuneus) for semantic processing, but not for phonological processing. The involvement of the bilateral parietal areas is related to retrieval of semantic information in adults (Binder and Desai, 2011). For children of age 9 years and older, the left IPL and angular gyrus have been reported to be specialized for semantic categorization tasks (Booth et al., 2007; Landi et al., 2010). Our study provided evidence of bilateral parietal involvement (including the supramarginal gyrus, angular gyrus and precuneus) for semantic processing not for phonological processing, suggesting specialization of bilateral parietal regions for semantic processing presents even in young children (5-7 years of age). In contrast, Weiss et al. did not observe parietal specialization for semantic categorization tasks using auditory stimuli for young children (5-6 years of age) (Weiss et al., 2018). They argued that parietal regions might be specialized for semantic processing only later in development. But our results are in line with another study that identified the bilateral right inferior parietal cortex during implicit processing of visually presented words (Turkeltaub et al., 2003). Thus, we proposed that the visual stimuli implemented in our study require high imageability for semantic decisionmaking, and bilateral parietal areas were thus recruited for visual semantic categorization tasks in young children.

### Implications on Theoretical Models of Reading

The DRC model is a particularly good example of the weak-phonological perspective that a direct lexical route takes precedence and is supported by a slow, secondary, and nonessential indirect phonological coding route (Coltheart et al., 2001). The neural pathways of the DRC model have been suggested to involve the dorsal and ventral pathways (Turkeltaub et al., 2003). The present study provided the neural bases of pre-reading skills and supports the DRC model. Our results indicated the involvement of dorsal pathway for phonological processing as evidenced by the functional specialization of sub-regions of the left IFG. Our study also provided evidence for the PDP model illustrated by common activation of brain regions related to phonology and semantics in young children. Binder et al. studied 24 healthy adults using word naming task and also supported the PDP model (Binder et al., 2005).

## Limitations

In total, 60 children completed the first behavioral testing session, but only 20 of 60 children were invited back to the second fMRI session (33%) based on their pre-reading skills, which might bias our sample. This is one of the limitations of the present study. We did not observe any significant brain-behavior correlation which can be attributed to the fact that all 19 children are good prereaders. The behavioral measures of their pre-reading skills had ceiling effects, which makes brain-behavior correlation hard to be significant and limits the generalization of our results. As we are interested in the neural bases of rhyming and semantics, children need to be able to complete tasks with above 60% accuracy. We could try to use lower-level cognitive tasks such as first-sound matching (Raschle et al., 2014; Yu et al., 2018) or letter identification (Yamada et al., 2011), but these tasks would be too simple for those who are 7 years old in our study. We also collected task-free resting-state fMRI data on our sample, but resting-state fMRI data could not provide task-specific information related to rhyming and semantics.

## Conclusions and Future Directions

The present study used multivariate approach to understand the neural bases of phonological and semantic processing in early childhood (5-7 years of age), which has not been reported previously. Our MVPA results suggested that a left lateralization for the indirect grapho-phonological route has already shaped in young children with good reading ability. Moreover, the lexico-semantic route also relies on the left hemisphere regions including the left IFG/STG/MTG/FG, and the additional recruitment of the bilateral parietal regions. Our MVPA results found the left opIFG specialization for the phonological processing, aligned with previous research in adults (Jobard et al., 2003) and school-age children (Bach et al., 2010). Our study was not designed to test the DRC and PDP models and thus our findings on the neural bases of phonological and semantic processing in early childhood did not conclusively support either model. Future studies with specific tasks targeting the two models are needed to examine the neural evidence for the two models. For future research, age-appropriate cognitive paradigms are required to identify pre-reading processes in even younger children (3-5 years of age). Moreover, within-subject design will help to determine the effects of modality (visual versus auditory) in neural specialization of phonological and semantic processes in early childhood.

## Supporting information

comments

## Author Contribution Statement

Mathur, A. created the task paradigms using E-prime 2, helped with data collection, analyzed the data, and drafted the manuscript with supports from Wang, Y.. Wang, Y. supervised the entire project and contributed to the conception, design, and data collection of the present study, and revising the manuscript for important intellectual content. Schultz, D. provided consultation on MVPA analysis.

## Acknowledgements

The authors thank the families for their participation. The authors also thank the assistance from undergraduate research assistants: Cristal Franco-Granados, Makayla Gill, Emily Ann Grybas, Meredith Konkol, Linneaa Nguyen, Grace Oh, Michelle Rohman, Fatima Sibaii, Thy Thy Trat Thai for helping with recruitment and data collection. This work was supported by funds from the Barkley Trust, Nebraska Tobacco Settlement Biomedical Research Development, College of Education and Human Sciences, and the Office of Research and Economic Development at University of Nebraska-Lincoln (UNL) and the Layman Fund (awarded to Wang, Y.) from the University of Nebraska Foundation. In addition, this work was supported by the National Institute of General Medical Sciences, 1U54GM115458, which funds the Great Plains IDEA-CTR Network. The content of this paper is solely the responsibility of the authors and does not necessarily represent the official views of the NIH. Some of the undergraduate research assistants were supported by the UNL Undergraduate Creative Activities and Research Experience (UCARE) program funded in part by gifts from the Pepsi Quasi Endowment and Union Bank & Trust.

## Acronyms

BOLD: blood-oxygen-level-dependent
CELF: Clinical Evaluation of Language Fundamentals
DRC: Dual Route Cascaded
FG: fusiform gyrus
IFG: inferior frontal gyrus
opIFG: opercular part of IFG
orIFG: orbital part of IFG
trIFG: triangular part of IFG
IPL: inferior parietal lobe
ITG: inferior temporal gyrus
LDA: linear discriminant analysis
LSA: Latent Semantic Analysis
MFG: mid-frontal gyrus
MOG: middle occipital gyrus
MTG: middle temporal gyrus
MVPA: multivariate pattern-analysis
PA: Phonological Awareness
PDP: Parallel Distributed Processing
RT: reaction time
SMA: supplementary motor area
SPL: superior parietal lobule
SFG: superior frontal gyrus
STG: superior temporal gyrus
TR: Repetition Time
VWFA: visual word form area
WRMT: Woodcock Reading Mastery Test

---

Supplementary Figure 1. Voxel-wise significant activation, within the whole brain anatomical mask for the contrast rhyming vs control. The color bar indicates test statistics (t-score) for the significant clusters identified using a voxel-wise threshold of q < 0.01 uncorrected, k >20.

