## supplementary table 1 for "Neural bases of phonological and semantic processing in early childhood"

Supplementary Table 1. Conjunction analysis results for Rhyming vs Control and Semantic vs Control.

| Region | L/R | BA | MNI Coordinates | | | Cluster Size | z-value | p-value* |
| --- | --- | --- | --- | --- | --- | --- | --- | --- |
|  |  |  | x | y | z |  |  |  |
| **Frontal** | | | | | | | | |
| IFG | L | 44/45/47 | -45 | 32 | 6 | 735 | 5.81 | <0.001 |
| SMA | L | 6 | -5 | 12 | 60` | 81 | 4.98 | 0.001 |

* p-value survived cluster-based (k > 20) FDR correction (q < 0.05).
