## Supplementary material for "Neural bases of phonological and semantic processing in early childhood": comments

UNIVERSITY OF Nebraska Lincoln
COLLEGE OF EDUCATION AND HUMAN SCIENCES
Department of Special Education and Communication Disorders
*Serving People with Special Needs*

Dr. Avantika Mathur, Postdoctoral Fellow
Department of Special Education and Communication Disorders (SECD)
Center for Brain, Biology and Behavior (CB3)
University of Nebraska-Lincoln (UNL)
 I Website: http://thewanglab.com

March 1, 2020

Dr. Paul Edison
Editor in Chief, Brain Connectivity


Manuscript ID BRAIN-2019-0728

Dear Dr. Edison,

Thank you for allowing us to revise our manuscript (ID: BRAIN-2019-0728) to be considered for publication in *Brain connectivity*. We much appreciate the time and efforts that you and the reviewers have dedicated to providing your valuable feedback on our manuscript. We are grateful to the reviewers for their insightful comments on our paper. We have been able to incorporate changes to reflect most of the suggestions provided by the reviewer. We have highlighted the revisions within the manuscript. Here are point-by-point responses to the reviewer's comments and concerns, which are highlighted in the red fonts. Each author listed on the submission made significantly contributions and the author list is complete. No fraudulent or manipulated data was used and all reported data and results are authentic and valid. No content of the submission has been plagiarized.

**Comments from Reviewer 1**

Comments to the Author

This fMRI study investigates the functional organization of phonological and semantic processing in 19 typically developing children between 5 and 7 years old. It is an important study contributing to our understanding of how neural networks underlying reading acquisition develop in beginning readers, especially given the lack of neuroimaging evidence in such a young age range. The combination of the univariate and multivariate analysis approaches provide evidence for functional specialization of the phonological (indirect) vs. semantic (direct) routes. The authors took great care for quality control during the behavioral and imaging data collection via the use of the mock scanner and sufficient task practice before scanning these young children. The analyses were rigorous. This dataset offers great potential for our understanding between children's reading skills and brain organization. However, the authors did not mention any brain-behavior relationship in this manuscript. In particular, I am curious about the implication of greater neural specialization in this age. In addition, the neural specialization vs. hemispheric specialization for reading was not well defined throughout the manuscript. More clarification is required for a better understanding of the results. I also suggest some additional analyses that might help authors to interpret their results.

We thank the reviewer for appreciating the efforts put forth in data collection and analysis for this study. Thank you for pointing out that clarity was missed in defining neural specialization vs. hemispheric specialization, we have taken care of the terms that we used throughout the manuscript and highlighted the changes in yellow. To be more precise, we changed the title and running head for this manuscript. We agree with the reviewer that we did not mention about brain-behavior correlation, which is one of the limitations of our study. The 20 children recruited in our study were good readers selected after screening 60 children on their reading ability. The behavioral measures had ceiling effects, which limits the generalization of the present work and makes brain-behavior correlation hard to be significant. Therefore, we described the challenge in the limitation section of the manuscript. By the reviewer's comments, we did additional analyses, including covariate analysis using accuracy scores as covariates of no interest and conjunction analysis. You can find details of the additional analyses in the relevant sections of our responses.

**Introduction:**

The theoretic overview for reading in adults is a little biased. It will be useful for readers to understand how the dual-route theory manifests during natural reading in adults. In addition, The dual-route theory for reading, albeit dominating the field, does not take learning into account. In the contrast, the parallel distributed processing model (Seidenberg & McClelland, 1989; Plaut, McClelland, Seidenberg, & Patterson, 1996), another prominent theory in the reading literature, produces outputs stimulating children's learning behavior. The authors should consider either to include this theory in the overview in the introduction or mention the implications of the results for this theory in the discussion.
We thank the reviewer for the suggestion of adding the parallel distributed processing (PDP) model. We have edited our introduction and added description of both theory models in the introduction (see yellow highlights on page 5 – 6). We have also discussed the implications of our results on the two theory models in the Discussion (see yellow highlights on page 21 – 22).

• Page 5: the gap of the literature in terms of "specialization" needs more clarification. Does greater functional specialization in a mature brain between the two pathways indicate better learning? Does the functional specialization mean hemispheric specialization on the left hemisphere or the dissociation between the two pathways? The discussion section of this paper suggests both aspects of the functional specialization has been examined in this paper. Please clarify.
We thank the reviewer for pointing out that clarity was missed in defining neural specialization vs. hemispheric specialization, we have addressed that throughout the manuscript (see highlighted changes).

**Methods:**

• Please indicate the voxel-level (cluster-defining) p-value for the univariate analyses.
The results for univariate analysis are reported at FDR corrected threshold of $q < 0.05$. We have added the peak level FDR-corrected values for each of the reported location in the univariate Tables 2 and 3.

Results:

• Please mark what the color scales indicate in Figure 3, Figure 4, and supplemental fig 1.

We thank the reviewer for the suggestion. For univariate analysis, Figure 3, the color bar indicates test statistics (t-score) for the significant clusters identified using a voxel-wise threshold of $q < 0.05$, FDR correction at a cluster threshold $k > 20$. For multivariate analysis, the color bar represents the test statistics (z-score) for the clusters with significant decoding accuracy. We have added the meaning of color scales in the figure legends.

• There is no direct comparison between the two tasks.

Thanks for pointing this out. The direct comparison between tasks did not reveal any significant differences at a corrected threshold. We have added the null results on page 19.

• Is the activation at the left IFG due to the task engagement or the activation of the language network? Because the control condition is easier than both the phonological and the semantic conditions, it is important to show that the condition contrasts was not due to task difficulty. suggest the authors also run an analysis to control for the accuracy difference.

We thank the reviewer for suggestion. We have re-analyzed the data using the task accuracy scores as covariate of no interest and updated the results. After controlling for accuracy, left IFG activation sustained at a corrected threshold (FDR, $q<0.05$) for both rhyming and semantic task, indicating the activation is due to activation of language network. The Figure 3 has been redone and results have been re-reported in tables 2 and 3 after running the covariate analysis.

• I suggest providing some quantifiable information about the neural specialization of the phonological vs. semantic task (e.g. a conjunction analysis), especially because the studies that directly compared brain activation patterns between the two tasks are extremely limited (as mentioned by the authors on page 5).

Thank you for this suggestion. We performed additional analysis and added the results of the conjunction analysis (in supplementary Figure 2 and supplementary Table1). The conjunction analysis performed between the two contrasts rhyme versus control and semantic versus control revealed significant overlapping activity within the left IFG and the left SMA at FDR-corrected threshold ($q < 0.05$) (page 16).

Discussion:

• Though it is helpful to comment on the converging evidence between univariate and multivariate analyses, it is also important to provide explanations and guidance for readers to digest the different results revealed by the analyses. For example: how to interpret the engagement of left IFG in both tasks revealed by the univariate analyses, but only in the semantic task revealed by the multivariate analyses?

We agree with the reviewer and have edited the discussion (see page 18 – 20). The major difference between the two approaches was observed in identifying the specialization of the sub-regions of the left IFG for phonological and semantic processing. Using univariate analysis approach, we did not observe any significant difference between the two tasks in IFG sub-regions, as subtraction analysis revealed no significant clusters between the two tasks and the conjunction analysis revealed an overlap in IFG sub-regions between the two tasks.

But, with multivariate analysis approach, we did observe a significant difference in the IFG sub-regions for the two tasks. Specifically, searchlight results for Rhyme vs Control revealed significant decoding accuracy in the cluster enclosing L IFG opercularis only, extending to pre/postcentral gyrus, cluster size 1380 voxels (refer to Table 4 and please see the attached cross-section image below, sup-Figure 1). On the other hand, searchlight results for Semantic vs Control revealed significant decoding accuracy in all the L IFG sub-regions (p. tri, p. op. and p. orb.), cluster size 1680 voxels (refer to Table 5 and please see the attached cross-section image below, sup-Figure 2).

The results included the Table 4 and Table 5 with the manuscript and discussion is modified in the manuscript [page 21 – 23].

sup-Figure 1. Cross – section image representing MVPA searchlight results for Rhyme vs Control.

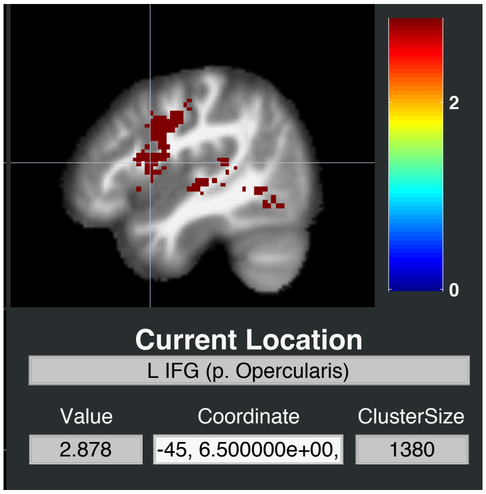


sup-Figure 2. Cross – section image representing MVPA searchlight results for Semantic vs Control.

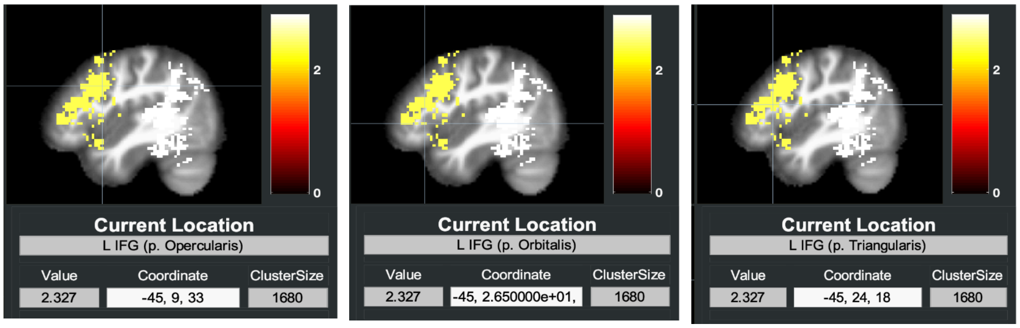
