## Supplementary figures and images for "Neural bases of phonological and semantic processing in early childhood"

### supplementary figure 1

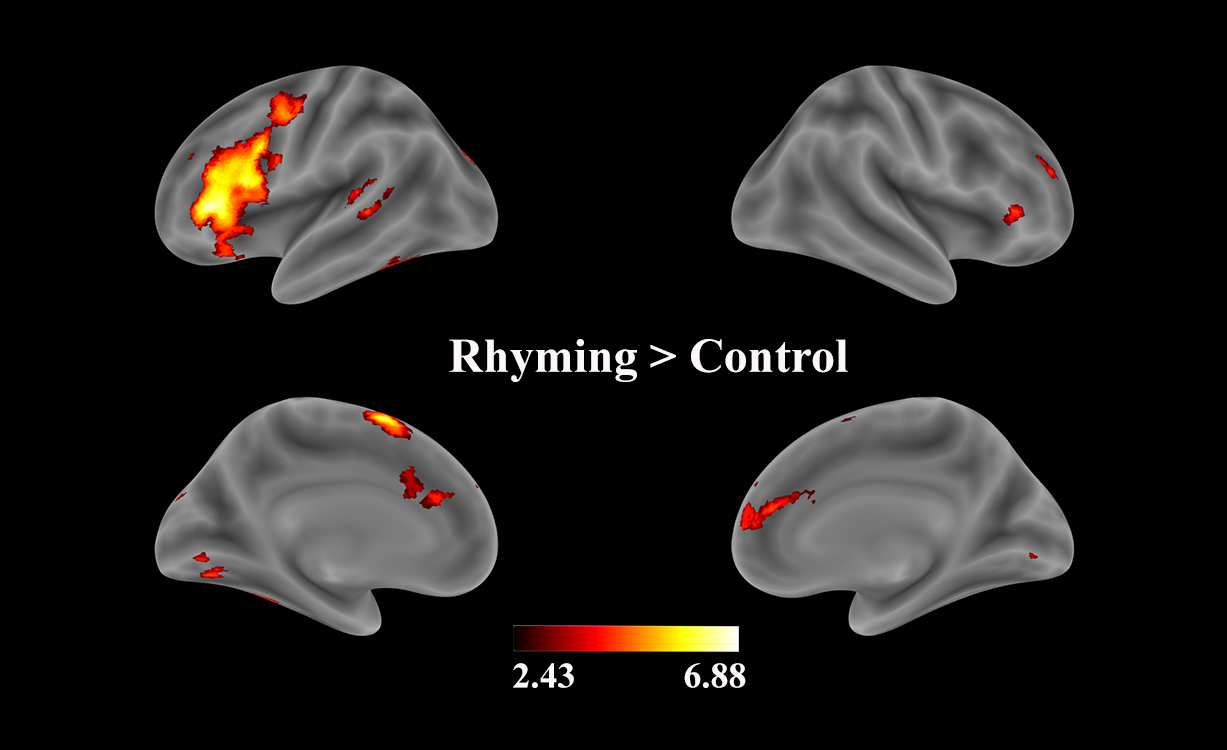

### supplementary figure 2

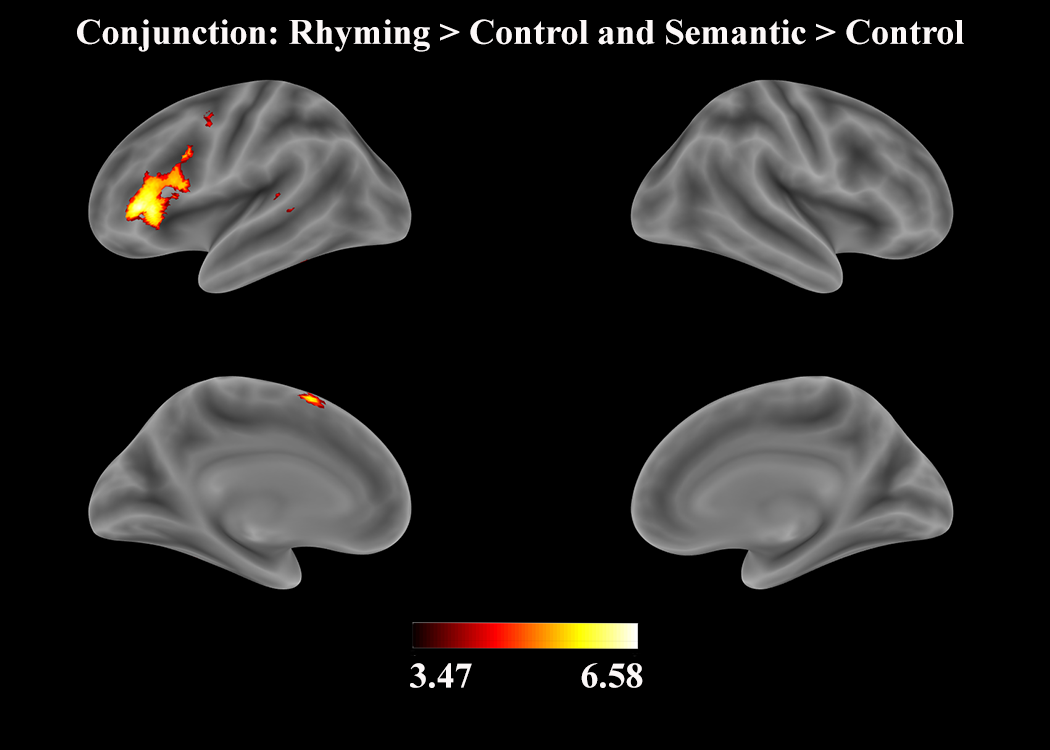
